# Inhibition of aryl hydrocarbon receptor signaling promotes the terminal differentiation of human erythrocytes

**DOI:** 10.1101/2021.06.08.447482

**Authors:** Yijin Chen, Yong Dong, Xulin Lu, Wanjing Li, Yimeng Zhang, Bin Mao, Xu Pan, Xiaohong Li, Ya Zhou, Quanming An, Fangxin Xie, Shihui Wang, Yuan Xue, Xinping Cai, Mowen Lai, Qiongxiu Zhou, Yan Yan, Ruohan Fu, Hong Wang, Tatsutoshi Nakahata, Xiuli An, Lihong Shi, Yonggang Zhang, Feng Ma

## Abstract

The aryl hydrocarbon receptor (AHR) plays an important role during mammalian embryo development. Inhibition of AHR signaling promotes the development of hematopoietic stem/progenitor cells. AHR also regulates the functional maturation of blood cells, such as T cells and megakaryocytes. However, little is known about the role of AHR modulation during the development of erythroid cells. In this study, we used the AHR antagonist StemRegenin 1 (SR1) and the AHR agonist 2, 3, 7, 8-tetrachlorodibenzo-*p-*dioxin (TCDD) during different stages of human erythropoiesis to elucidate the function of AHR. We found that antagonizing AHR signaling improved the production of human embryonic stem cell (hESC)-derived erythrocytes and enhanced erythroid terminal differentiation. RNA-sequencing showed that SR1 treatment of proerythroblasts upregulated the expression of erythrocyte differentiation-related genes and downregulated actin organization-associated genes. We found that SR1 promoted F-actin remodeling in terminally differentiated erythrocytes, favoring the maturation of the cytoskeleton and enucleation. We demonstrated that the effects of AHR inhibition on erythroid maturation resulted from an increase in F-actin remodeling. Our findings help uncover the mechanism for AHR-mediated human erythroid cell differentiation. We also provide a new approach toward the large-scale production of functionally mature hPSC-derived erythrocytes for use in translational applications.

## Introduction

The aryl hydrocarbon receptor (AHR) is a member of the basic-helix-loop-helix Per-ARNT-SIM family of transcription factors, which play important roles in various physiological processes^1^. AHR can be activated by endogenous or exogenous ligands, and is involved in the regulation of inflammation^2–4^, cell differentiation^5, 6^, apoptosis^7^, cancer progression^8^, and neural function^9^. Several studies indicate that AHR has an essential role in hematopoiesis^10–14^. AHR^-/-^ mice show a marked increase in the number of hematopoietic stem cells (HSC) in the bone marrow and an increased propensity to develop lymphomas^15–17^. Cultures of human pluripotent stem cells (hPSC) treated with the AHR antagonist StemRegenin 1 (SR1), or in which the *AHR* gene was deleted, have increased levels of CD34^+^CD45^+^ hematopoietic progenitor cells (HPCs); there was also a significant increase in the number of colony-forming-unit erythroid (CFU-E) and colony-forming-unit macrophages (CFU-M)^11^. Moreover, recent research shows that AHR is involved in the determination of cell fate. Activation of AHR promotes cellular differentiation toward the formation of monocyte-derived dendritic cells by inhibiting the formation of macrophages^18^. Furthermore, AHR plays an important role in the differentiation of B cells^19, 20^, natural killer cells^21^ and megakaryocytes^22, 23^.

Following commitment to the erythroid lineage, HSC sequentially give rise to common myeloid progenitor, megakaryocyte-erythrocyte progenitor (MEP), burst-forming unit-erythroid (BFU-E) and CFU-E cells^24^. Subsequently, erythroid precursors enter terminal differentiation, sequentially produce morphologically recognizable proerythroblasts; basophilic, polychromatic, and orthochromatic erythroblasts; reticulocytes; and finally, they mature into red blood cells. During maturation, erythrocytes undergo serial changes during which the cell and the nucleus decrease in size, hemoglobin is synthesized, membrane proteins are reorganized, chromatin condensation is regulated, and finally, cells become enucleated^25, 26^. The expression of AHR is upregulated in HSCs and MEP cells, but gradually downregulated during erythroid differentiation^27^. This finding suggests that the differential modulation of AHR is important during the process of erythrocyte maturation. In MEP cells, chronic AHR activation causes differentiation of erythroid cells, and repression of AHR activity leads to megakaryocyte specification^28^. However, several reports show that exposure of zebrafish^29, 30^ and human ^31^ erythroblasts to the AHR agonist 2, 3, 7, 8-tetrachlorodibenzo-p-dioxin (TCDD) damaged the deformation ability of erythroblasts and decreased hemoglobin synthesis. Thus, the role of AHR in the development and differentiation of erythrocytes is still ambiguous.

Remodeling of F-actin is essential for nuclear extrusion during the terminal differentiation of erythroid cells^32^. The expression of actin reaches a peak in proerythroblasts, and F-actin forms a ring structure underneath the cytoplasmic membrane. During terminal differentiation, F-actin rearranges and forms a contractile ring between the extruding nucleus and the incipient reticulocyte, which benefits erythroblast enucleation^33–35^. F-actin remodeling is regulated by many factors, including AHR, which regulates the expression of actin-related genes to modulate cellular shape and function^36–39^. AHR activation by TCDD impairs the deformation ability of erythroid cells, indicating that AHR may have a role in the development of the erythroid cytoskeleton.

In this study, by an efficient co-culture system we established to inducing hematopoietic and erythrocyte differentiation from hPSCs^40–42^, we used the AHR antagonist SR1 to show that inhibiting AHR signaling promotes erythroid terminal differentiation. Particularly, SR1 promotes actin to be localized to one side of the cell near the nucleus, showing that AHR signaling regulates cytoskeletal remolding in mature erythrocytes. Furthermore, addition of SR1 to cultures of hESC-derived erythrocytes during terminal differentiation increases the number of enucleated erythroid cells that can be harvested. These findings increase our knowledge of AHR function during erythroid cell differentiation, and provide a possible way to enhance the production and maturation of hESC-derived mature erythrocytes.

## Results

### SR1 increases erythroid cell production from hESC

We have described previously an efficient system of producing HSPC from hESC by co-culturing with aorta-gonad-mesonephros (AGM)-S3^40, 41, 43^. In this system, hematopoietic endothelial cells (HEC; CD34^+^KDR^+^CD43^-^GPA^-^) and hematopoietic progenitor cells (HPC; CD34^+^CD43^+^CD45^-^GPA^-^) can be detected on co-D6 and co-D10^40^. In addition, HSPC (CD34^+^CD45^+^) emerge at co-D10, and reach a peak at co-D14. Early erythroblasts (CD34^+^GPA^+^) can also be observed before the emergence of HSPC^41^.

In the current study, SR1 was continuously added to the co-culture system from co-D6, and total cells were harvested every 2 days until co-D14 for fluorescence-activated cell sorting analysis and the colony-forming-unit assay (Figure 1A). As previously reported^11^, SR1 treatment significantly increased the number of HPC and HSPC compared with the DMSO-treated control (Figure 1B and C). Interestingly, SR1-mediated enhancement of HSPC production was accompanied by an increase in the number of erythroid (GPA^+^CD45^-^) and myeloid (CD45^+^GPA^-^CD34^-^) cells. Of note, SR1 significantly increased the number of early erythroblasts but not the number of myeloid cells at co-D8, when HSPCs have not emerged. Similar results were observed in the colony-forming-unit assay (Figure 1D). The production of granulocytic-erythroid-megakaryocytic-macrophagic (GEMM) colony-forming-unit was higher in SR1-treated cells than in control cells. In addition, SR1 treatment increased the production of other types of colonies. Interestingly, SR1 treatment caused a significant increase in the formation of CFU-E colonies form co-D8, whereas BFUE and myeloid (granulocyte or macrophage or granulocyte/macrophage, G/M/GM) colonies were increased at co-D10 when HSPCs emerged. These data indicate that AHR may have a role in erythropoiesis.

**Figure 1.**
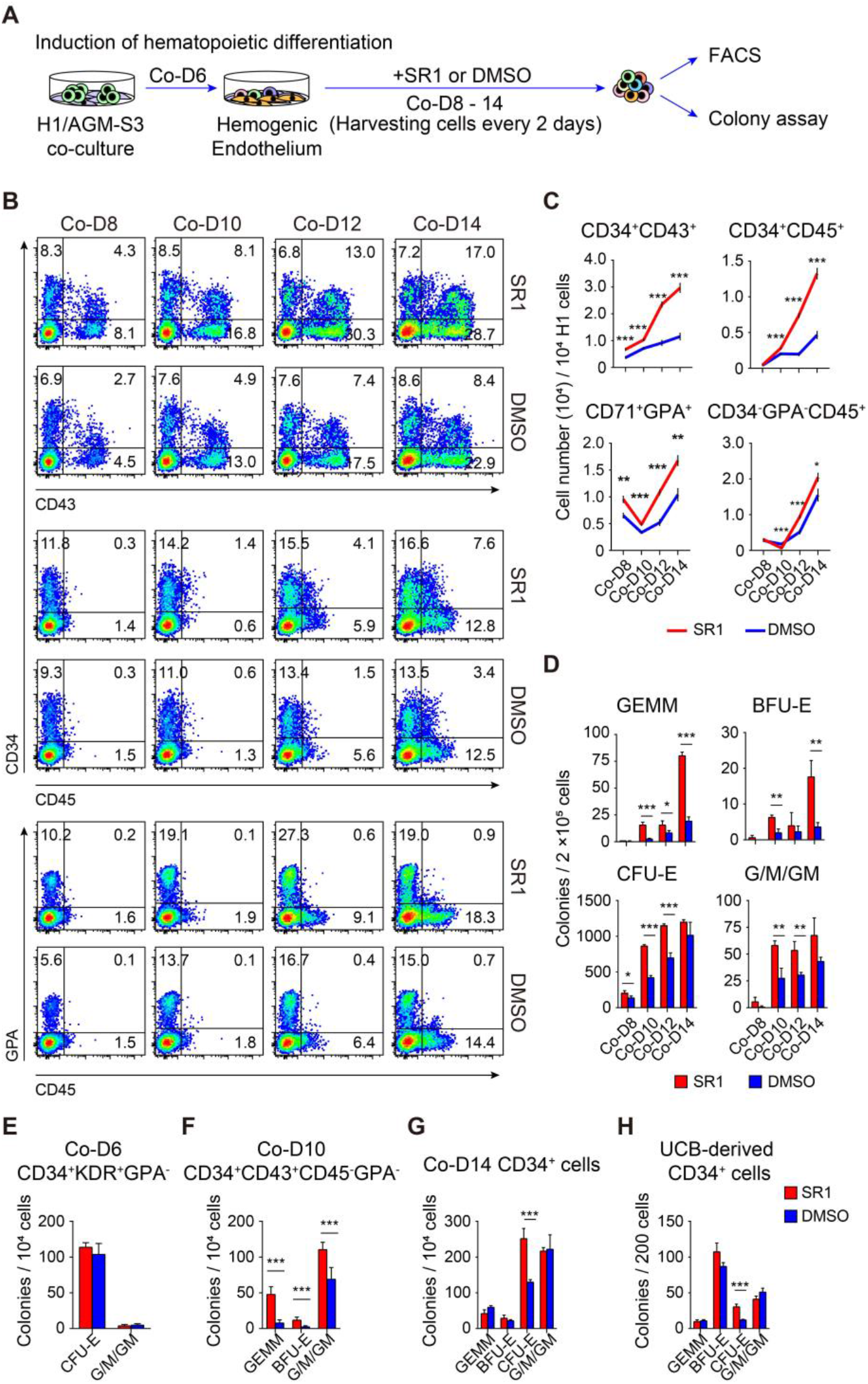
StemRegenin 1 (SR1) increases erythroid cell production from human embryo stem cell (hESC). (A) A schematic of the strategy used to analyze the effect of SR1 on hematopoiesis. SR1 was continuously added from D6 onwards during H1/AGM-S3 co-culture (co-D6), and total cells were harvested every 2 days for flow cytometry analysis and the colony-forming-unit assay. DMSO was added as the vehicle control. Flow cytometry analysis (B) and absolute cell number (C) of hematopoietic cells (CD34^+^CD43^+^), hematopoietic stem/progenitor cells (CD34^+^CD45^+^), erythroid cells (CD71^+^GPA^+^) and myeloid cells (CD34^-^GPA^-^CD45^+^) treated with or without SR1 from co-D8-14. (D) The clonogenic capacity of co-cultured cells treated with or without SR1 from co-D8-14. The number of granulocytic-erythroid-megakaryocytic-macrophagic (GEMM), burst-forming unit-erythroid (BFU-E), colony-forming-unit erythroid (CFU-E) and granulocyte or macrophage or granulocyte/macrophage (G/M/GM) colonies are shown. (E-G) Isolated co-D6 hematopoietic endothelial cells (CD34^+^KDR^+^GPA^-^) and co-D10 hematopoietic progenitor cells (CD34^+^CD43^+^CD45^-^GPA^-^) were replanted on AGM-S3 that included hematopoietic differentiation medium, and co-D14- and umbilical cord blood (UCB)-derived CD34+ cells were cultured in erythroid differentiation medium, and the clonogenic capacity of these cells treated with or without SR1 at D3 was measured. The number of colonies derived from hematopoietic endothelial cells (E), hematopoietic cells (F), co-D14 (G) or UCB-derived (H) CD34+ cells. All values are mean ± SD, N = 3; * p < 0.05, ** p < 0.01, *** p < 0.001.

To investigate whether AHR can influence erythroid development, sorted co-D6 HEC, co-D10 HPC and co-D14 CD34^+^ cells were exposed to SR1 for 3 days and used in the colony-forming-unit assay. More HPCs were generated from co-D6 HEC (Figure S1A and B), but there was no significant difference in HEC-derived CFU-E colonies between SR1- and DMSO-treated cells (Figure 1E). We also found that more HSPCs were generated from SR1-treated HPC, together with more GEMM-, BFU-E- and G/M/GM-producing colonies (Figure 1F; Figure S1C and B). However, we did not observe more GEMM, BFU-E and G/M/GM colonies in SR1-treated co-D14 CD34^+^ cells, but CFU-E colony formation increased during erythroid differentiation (Figure 1G). Although SR1 treatment did not significantly alter the number of erythroid cells (CD71^+^GPA^+^) at D3, a higher percentage of this cell type was detected in SR1-treated cultures compared with the DMSO-treated control (Figure S1E and F). Similar results were observed in UCB-derived CD34^+^ cells (Figure 1H). Although chronic SR1 treatment generated fewer cells, the percentage of erythroid cells was always higher in SR1-treated cultures than in DMSO-treated cultures (Figure S1G and H). Collectively, these data suggest that AHR antagonism has a positive effect on erythroid differentiation rather than generation from hematopoietic progenitors.

### AHR antagonism promotes erythroid terminal differentiation

We wondered if AHR inhibition increases CD71^+^GPA^+^ cells and CFU-E colonies by acting on BFU-E cells. To test this hypothesis, UCB-derived BFU-E cells isolated by the expression of the surface marker CD34^+^CD71^+^GPA^-^CD123^-^^24^ at D4 were continuously exposed to SR1 or DMSO. At D1, SR1 treatment did not affect the formation of CFU-E colonies compared with DMSO-treated cells, but at D2 and D3 SR1 treatment produced more CFU-E colonies (Figure 2A). Moreover, SR1 treatment significantly increased the number of erythroid cells (Figure 2B and S2A). These results suggest that AHR antagonism may increase the differentiation of CFU-E cells.

**Figure 2.**
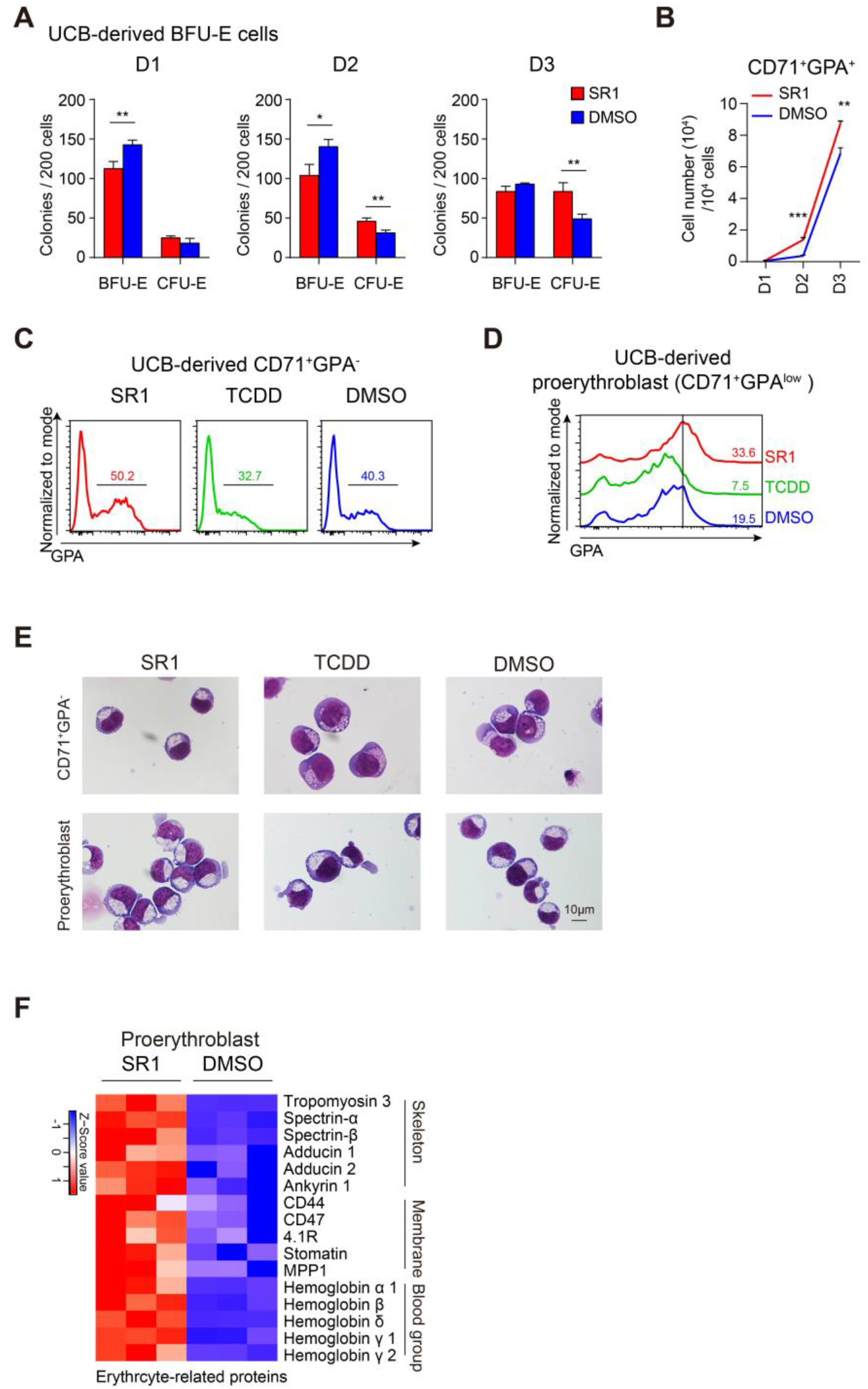
AHR antagonism promotes erythroid terminal differentiation. (A-B) Sorted UCB-derived BFU-E cells were treated continuously with or without SR1, and total cells were collected every day for flow cytometry analysis and the colony-forming-unit assay. The number of colonies (A) ant the number of CD71^+^GPA^+^ erythroid cells (B) at days 1, 2 and 3. (C, D) Isolated CFU-E enriched cells (CD71^+^GPA^-^) and proerythroblasts (CD71^+^GPA^low^) were treated with SR1, TCDD or DMSO. (C) Flow cytometry analysis of GPA^+^ cells generated from CD71^+^GPA^-^ cells at D3. (D) Flow cytometry analysis of GPA expression in proerythroblasts. (E) Representative images of May-Grunwald Giemsa (MGG)-stained cells; scale bar, 10μm. Isolated UCB-derived proerythroblasts were treated with or without SR1, and proteomic analysis of the total cells was performed at D3. (F) Heatmap representing the expression of erythroid-related proteins including skeletal, membrane, and blood-group proteins; columns represent the indicated replicates of each population; the colored bar shows row-standardized Z-scores. All values are mean ± SD, N = 3; * p < 0.05, ** p < 0.01, *** p < 0.001.

To further assess the role of AHR during CFU-E cells differentiation, D7 CFU-E-enriched cells (CD71^+^GPA^-^^43^) and proerythroblasts (CD71^+^GPA^low^) were continuously exposed to SR1, TCDD, or DMSO. Results showed that more GPA^+^ cells were generated from cultures of 71^+^GPA^-^ cells (Figure 2C), and the expression of GPA was higher in proerythroblasts (Figure 2D) under SR1-treated conditions at D3, TCDD significantly inhibited these processes. In addition, SR1 or TCDD treatment did not alter the number of CD71^+^GPA^-^ cells and proerythroblasts compared with DMSO treatment, although cultures treated with TCDD had a higher number of CD71^+^GPA^-^ cells than cultures treated with SR1 at D3 (Figure S2B and C). There was no significant difference in the level of apoptosis or changes in the cell cycle in cells derived from SR1-, TCDD- or DMSO-treated CD71^+^GPA^-^ cells or proerythroblasts (Figure S2D and E). SR1-treated CD71^+^GPA^-^ cells and proerythroblasts had a more mature shape at D3, whereas the shape of TCDD-treated cells reflected a more immature morphology (Figure 2E). Furthermore, proteomic analysis showed that the expression of skeleton proteins (tropomyosin 3, spectrin, adducin and ankyrin), membrane proteins (CD44, CD47, 4.1R, stomatin and MPP1), and blood-group proteins (hemoglobin α and β) was higher in SR1-treated proerythroblasts than in DMSO-treated proerythroblasts (Figure 2F). To test whether other agonists or antagonists of AHR had similar effects on erythroid terminal differentiation, UCB-derived proerythroblasts were exposed to 2-(1’H-indole-3’-carbonyl)thiazole- 4-carboxylic acid methyl ester (ITE; an endogenous agonist), 6-formylindolo[3,2-b]-carbazole (FICZ; an endogenous agonist) or N-(2-(1H-indol-3-yl)ethyl)-9-isopropyl-2-(5-methylpyridin-3-yl)-9H-purin-6-amin e (GNF351; an antagonist) for 3 days. GNF351 promote terminal differentiation, but ITE and FICZ prevented terminal differentiation (Figure S2F).

Next, we wondered if AHR had similar effects on H1-derived erythroid cells. Co-D14 CD34^+^ cell-derived CD71^+^GPA^-^ cells and proerythroblasts were sorted at D7 of erythroid differentiation. Similar trends to those seen in UCB-derived cells were observed in H1-derived cells (Figure S3A-C). Moreover, the production of CFU-E colonies was higher in both SR1-treated CD71^+^GPA^-^ cells and proerythroblasts than DMSO-treated cells (Figure S3D). In addition, SR1 enhanced the differentiation of H1-derived early erythroblasts and the ability to form CFU-E colonies (Figure S3E and F). Taken together, these results demonstrate that AHR inhibition promotes the terminal differentiation of human erythroid cells, whereas AHR activation arrests this process.

### Inhibition of AHR influences heme- and actin-related gene expression during terminal erythroid differentiation

To determine the cellular processes by which AHR antagonism enhances terminal differentiation, H1- or UCB-derived proerythroblasts treated for 3 days with SR1 were collected for RNA-seq analysis. SR1 treatment of UCB-derived cells upregulated the expression of 460 out of 1282 genes and downregulated the expression of 822 genes (Figure 3A). Similar patterns were observed in H1-derived cells (expression of 1365 genes was upregulated and expression of 1736 genes was downregulated). The expression of AHR-regulated genes such as cytochrome P450 1B1 (*CYP1B1*) and *AHRR* was markedly downregulated in SR1-treated cells (Figure S4A). To find upregulated and downregulated genes common to SR1-treated H1- and UCB-derived erythroid cells, we overlapped transcriptomic profiles from these two cell types. We found that the expression of 162 upregulated and 440 downregulated genes was common to both cell types. Next, gene ontology enrichment analysis was performed on these genes. Results showed that upregulated genes were highly associated with erythrocyte differentiation and homeostasis, as well as heme metabolic processes (Figure 3B). Genes involved in the erythrocyte differentiation processes were hemoglobin synthesis-related genes (*AHSP*, *ALAS2*, *TMEM14C* and *SOX6*), membrane protein-related genes (*SLC4A1*, *EPB42*), and erythropoietin signal-related genes (*JAK2*, *MFHAS1*; Figure 3C). 440 SR1-downregulated genes were enriched in actin-related processes, such as actin filament organization and regulation of small GTPase- mediated signal transduction (Figure 3D). Genes involved in the actin-related processes including *ARHGAP18*, *RAC2* and *SAMD14* (Figure 4SC). Furthermore, gene set enrichment analysis showed that SR1 treatment downregulated several actin cytoskeleton-associated genes (Figure 3E). Together, these data show that AHR inhibition upregulates the expression of erythrocyte differentiation- and heme metabolism-related genes, and downregulates the expression of actin-associated genes. This finding ensures that AHR inhibition promotes terminal erythroid differentiation, and suggests that AHR can modulate the function of the actin cytoskeleton during this process.

**Figure 3.**
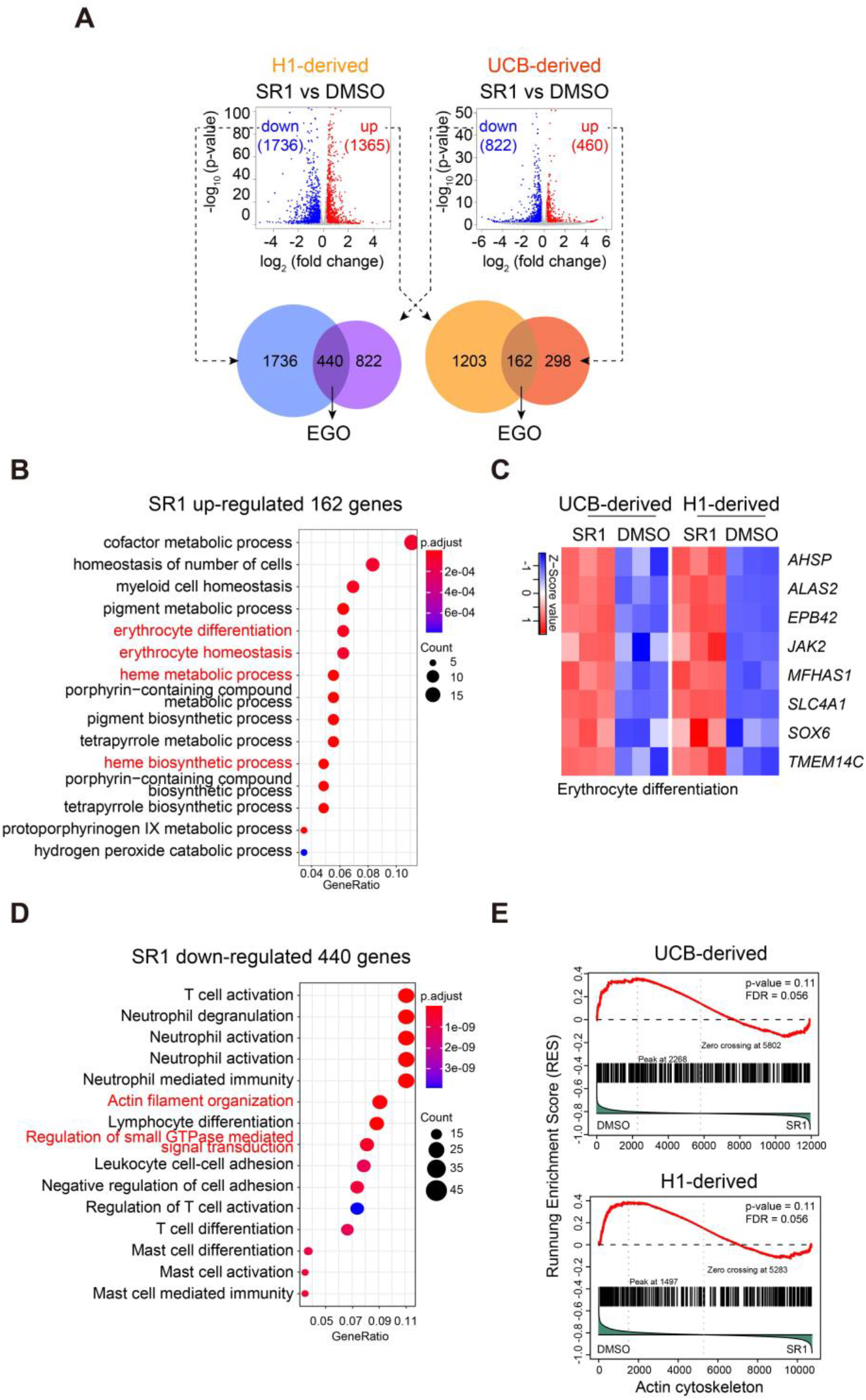
Inhibition of AHR influences heme- and actin-related gene expression during terminal erythroid differentiation. H1- or UCB-derived proerythroblasts were treated with or without SR1 for 3 days; total cells were harvested, and RNA sequencing was performed. (A) The correlation between RNA expression levels in H1- and UCB-derived proerythroblasts. Detected genes were filtered by p-value <0.05 and fold change >1.2. Differentially expressed RNA transcripts (red, upregulated; blue, downregulated) were assessed for overlap. Gene ontology enrichment analysis (EGO) of genes upregulated in both H1- and UCB-derived cells was performed. EGO analysis of the 162 co-upregulated (B) and 440 co-downregulated (D) genes using the clusterProfilter R package: each symbol represents a GO term (noted in the plot); the color indicates the adjusted p-value. (D) GSEA of UCB- or H1-derived actin cytoskeleton genes. (C) Heatmap analysis of erythrocyte differentiation-related genes identified from the results of EGO analysis. Columns represent the indicated replicates of each population; the color bar shows row-standardized Z-scores. (E) Gene set enrichment analysis (GSEA) of UCB- or H1-derived actin cytoskeleton genes. Pathways from the dataset of c2.all.v5.2.symbols downloaded from the GSEA website are shown.

### AHR inhibition enhances actin remodeling during erythroid maturation

Next, we observed actin expression during SR1 treatment. RNA-seq results showed that *ACTB* expression was decreased in SR1 treatment proerythroblasts (Figure 4A). Western blot results showed that SR1-treatment decreased β-actin expression in proerythroblasts (Figure 4B). Phalloidin was used as a marker of F-actin expression. We found that the amount of F-actin gradually decreases during the differentiation of proerythroblasts. Moreover, cells with lower F-actin expression had actin rearrangement at D3 (red arrow), and polarized F-actin was formed, which localized at one side of the nucleus at D5 (white arrow; Figure 4C). There was a significant increase in F-actin remodeling at D3, and actin polarization was more obvious on D5 in SR1-treated cells, TCDD prevented F-actin remodeling and actin polarization (Figure 4C). To further examine the function of AHR on actin remodeling, two actin-disrupting drugs, cytochalasin-D (CytoD) and jasplakinolide, were used. Surprisingly, CytoD, which blocks polymerization and elongation of F-actin, did not influence the differentiation of proerythroblasts, but reversed the action of SR1 on these cells (Figure 4D). Jasplakinolide, which stabilizes F-actin, produced similar results on SR1-treated proerythroblasts (Figure 4E). Similar results to those produced by CytoD, which reverse the effect of SR1 on proerythroblasts differentiation, were observed in GNF351-treated UCB-derived cells (Figure 4F) and SR1-treated H1-derived proerythroblasts (Figure 4G). These data confirmed that AHR antagonism accelerates F-actin remodeling and that disrupting F-actin remodeling blocks the promoting effect of AHR antagonism on erythroid terminal differentiation.

**Figure 4.**
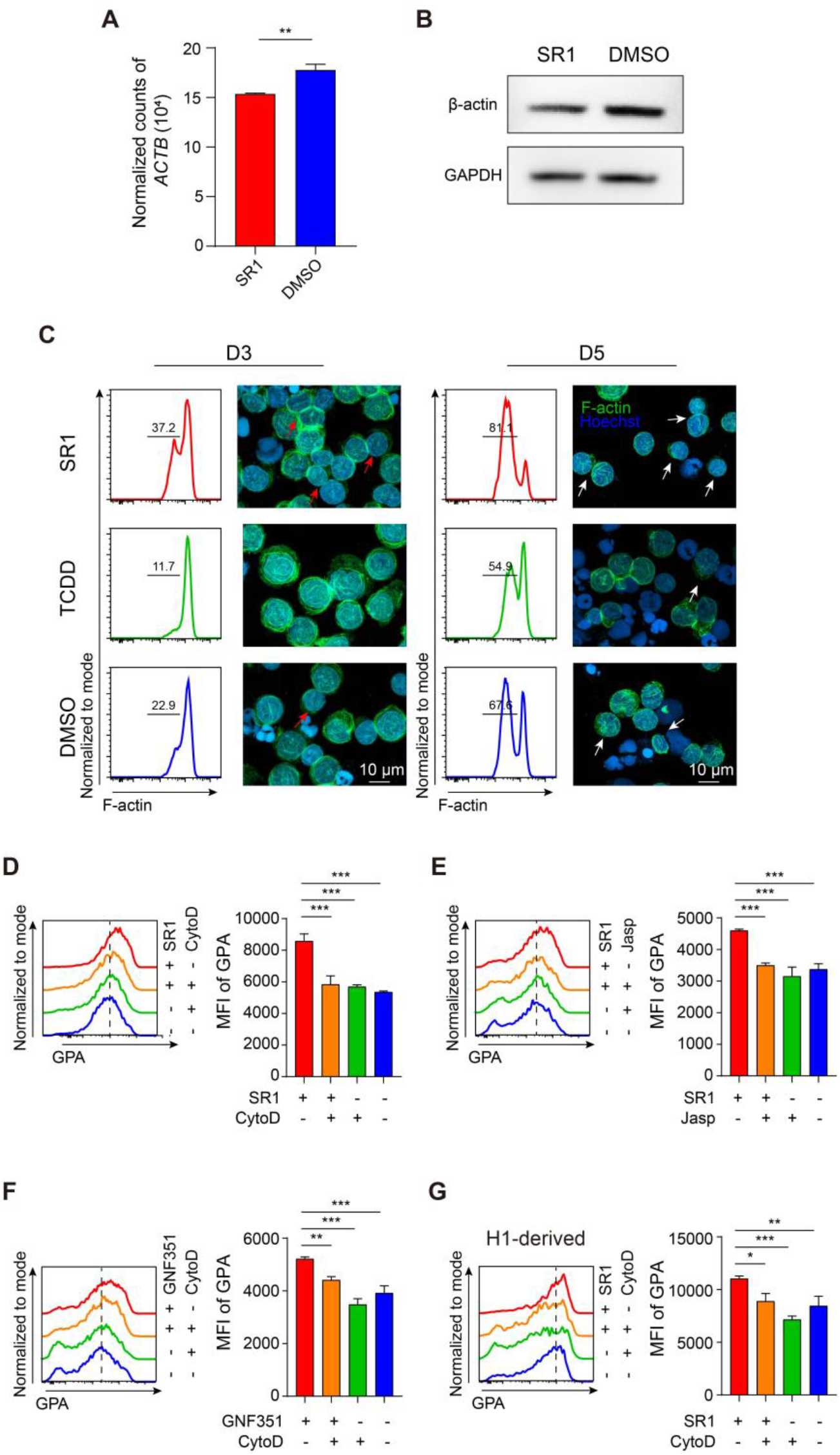
AHR inhibition enhances actin remodeling during erythroid maturation. The expression of β-actin in SR1-treated UCB-derived proerythroblasts was detected by RNA-seq (A) and western blot analysis (B) at D3. F-actin was detected in UCB-derived proerythroblasts continuously treated with SR1, TCDD, or DMSO at D3 and 5. (C) The proportion of F-actin was analyzed by flow cytometry, and structure of F-actin was observed by immunofluorescence staining; phalloidin (green staining) represents F-actin, and nuclei are stained blue; the red arrow indicates actin remodeling, the white arrow indicates polarized actin; scale bar, 20 μm. UCB-derived proerythroblasts were treated with or without the actin-disrupting drugs, cytochalasin-D (CytoD) or jasplakinolide (Jasp) under SR1- or DMSO-treated conditions. GPA expression in CytoD-treated (D) and Jasp-treated (E) cells was analyzed by flow cytometry at D3, and the mean fluorescence intensity (MFI) of GPA in these cells was calculated. UCB-derived CD71^+^GPA^low^ cells were treated with or without CytoD under GNF351- or DMSO-treated conditions. (F) GPA expression was analyzed by flow cytometry and MFI of GPA was calculated at D3. H1-derived proerythroblasts were treated with or without CytoD under SR1- or DMSO-treated conditions. (G) GPA expression was analyzed by flow cytometry and MFI of GPA was calculated at D3. All values are mean ± SD, N = 3; * p < 0.05, ** p < 0.01, *** p < 0.001.

### SR1 promotes the functional maturation of human erythroid cells

Since SR1 promotes terminal erythroid differentiation, we assessed whether SR1 modulated the maturation of human erythroid cells. Co-D14 or UCB-derived CD34^+^ cells were collected and re-cultured in erythroid differentiation medium; SR1 was added at D7 (Figure 5A). First, we examined the expression of erythroid-related proteins on co-D14-derived erythroid cells at day 12. Under SR1-treatment conditions, the expression of Band3, CD47 and hemoglobin-β was higher, but CD49d expression was lower (Figure S5A and B). However, the expression of CD29 and CD44 did not change (Figure S5A). At D21, the concentration of hemoglobin was higher in SR1-treated erythroid cells than in DMSO-treated cells; 45 g pre 10^12^ in SR1-treated cells and 40 g per 10^12^ in DMSO-treated cells (Figure 5B and C). Moreover, the number of live cells was about 2-fold higher in SR1-treated cultures than in the DMSO-treated control (Figure 5D). We next examined the enucleation rate of these cells. The results showed that SR1 increased the percentage of enucleated erythroid cells by about 2-fold (Figure 5E and F). In addition, we measured the deformation of these cells; SR1-treated cells had a higher deformation index under conditions of 1000 s^-1^ shear stress (Figure 5I). Similar results were obtained in UCB-derived erythroid cells (Figure 5C-D, and G-I). These data further support the notion that AHR inhibition promotes erythroid terminal differentiation and function maturation. Furthermore, we showed that H1-derived mature erythroid cells can be obtained by adding AHR antagonists to cell cultures during terminal differentiation.

**Figure 5.**
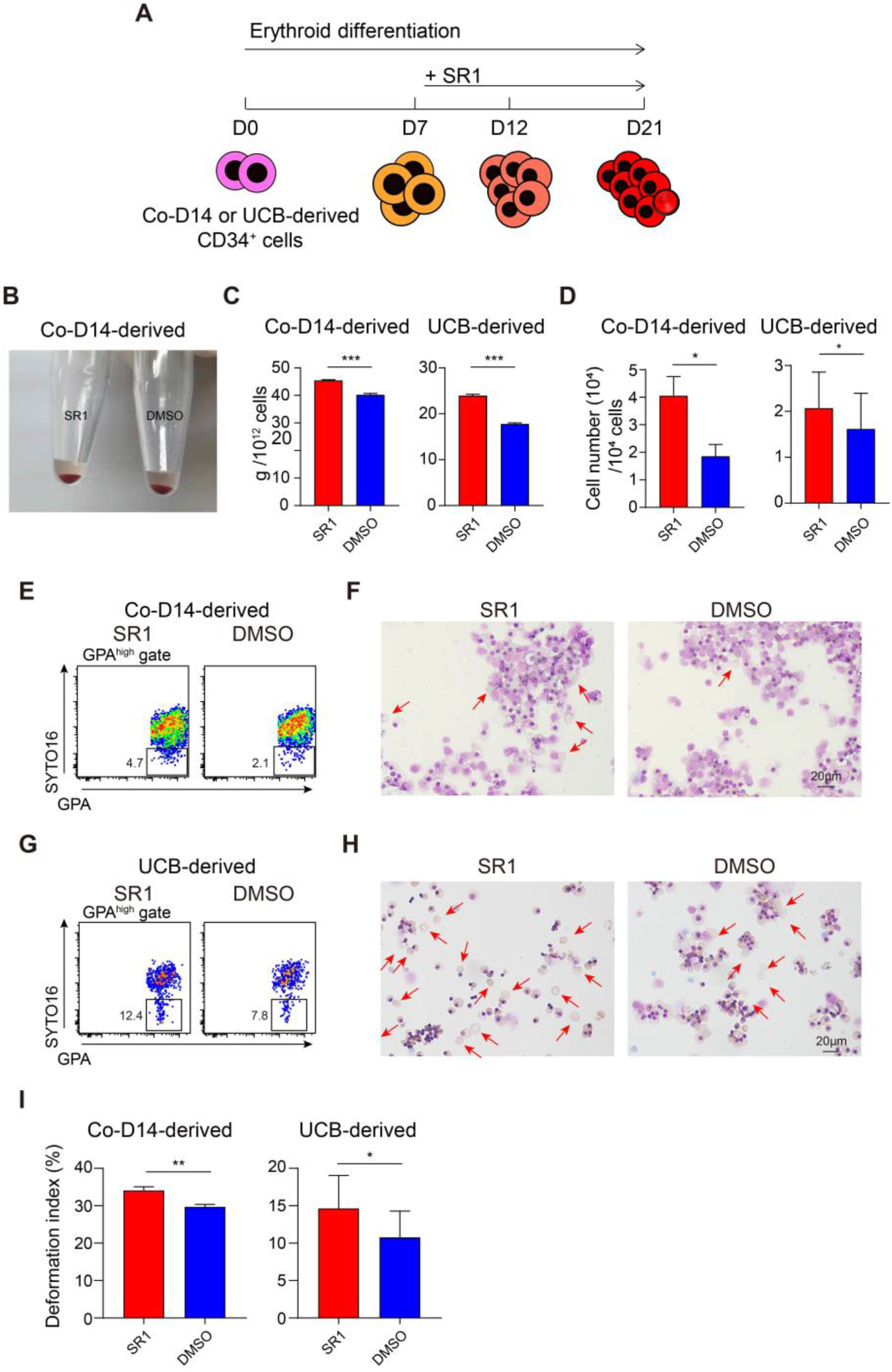
SR1 promotes the functional maturation of human erythroid cells. (A) A schematic of the strategy used to produce functional maturation of human erythroid cells. SR1 was added on D7, and erythroid cells were collected at day 21. (B) Images show co-D14-derived erythroid cells harvested at D21. Hemoglobin concentration (C) and the number of live cells (D) of Co-D14- and UCB-derived erythroid cells were calculated at D21. The enucleation rate of Co-D14- (E) and UCB-derived (G) erythroid cells was analyzed by flow cytometry, and the morphology of Co-D14- (F) and UCB-derived (H) cells was observed by MGG staining at D21; representative images of erythroid cells are shown. The red arrow indicates enucleated erythrocytes; scale bar, 20 μm. (G) Deformability of Co-D14- and UCB-derived erythroid cells was detected at D21. All values are mean ± SD, N = 3; * p < 0.05, ** p < 0.01, *** p < 0.001.

## Discussion

AHR signaling plays profound roles in biological events, particularly in the development and maintenance of HSCs^12, 45^, and the functional maturation of immune cells^6, 11, 18, 46^ and megakaryocytes^27^, but little is known about how AHR regulates erythroid differentiation. In our present study, through a step-by-step examination of the effects of AHR on the differentiation of erythrocytes, we found that AHR plays an important role in erythroid terminal differentiation.

Many reports show that AHR impacts the expansion and differentiation of HSPC^10, 12, 45^, and that AHR antagonists such as SR1 and GNF351 promote the expansion and maintenance of HSPC^12, 47–49^. In transgenic experimental models, Ahr^-/-^ mice showed an increase on HSPCs in bone marrow^15^, and CFU-E colony formation is increased in AHR^-/-^ hESC^11^. Moreover, inhibition of AHR activity increased the differentiation of hematopoietic cells from hPSC^11^. These studies are in agreement with our current study, in which AHR antagonists promoted the development of hESC-derived HSPC in AGM-S3 co-culture. Particularly, by using SR1 to antagonize AHR signaling at several time-points, we observed an increase in the production of erythroid cells both from HSPC-independent and -dependent pathways during early human hematopoiesis (Figure 1). These data suggest that AHR has an important role in erythropoiesis. We did not observe an increase in the number of CFU-E colonies when HECs (D6 in co-culture) were exposed to SR1 for 3 days (Figure 1E). However, hematopoietic colonies containing erythroid cells (mixed colonies and BFU-E) that originated from HPC at comparatively late stages (D10 in co-culture) were increased, along with the formation of other myeloid colonies (Figure 1F). Furthermore, when co-D14 or UCB-derived CD34^+^ cells were treated with SR1 for 3 days, only the formation of CFU-E colonies was increased in secondary semisolid cultures (Figure 1G and H). These results showed that inhibition of AHR does not directly influence the development of erythroid cells from multipotent hematopoietic progenitors, but suggest that AHR inhibition might affect the late erythrocyte differentiation stage.

Commitment to an erythroid lineage is typically marked by the appearance of BFU-E cells, defined as CD34^+^CD71^low^GPA^-^CD36^-^CD123^-^^24^. In our current study, we found that SR1 mostly exerts its effect at the BFU-E stage to promote CFU-E production and the further maturation of erythrocytes. Activation of AHR can generate erythroid progenitors from hiPSC-derived erythroid-megakaryocyte precursors^27^. Consistent with this report, we observed a moderate decrease in erythrocyte production from UCB-CD34^+^ cells when SR1 was continuously added to the culture (Figure S1H). However, when SR1 treatment was pulsed within 3 days, purified UCB-BFU-E cells produced more CFU-E colonies in semisolid culture and produced CD71^+^GPA^+^ cells in suspension culture (Figure 2A and B), and UCB-CFU-E cells produced more CD71^+^GPA^+^ cells as well (Figure 2C). Furthermore, antagonizing AHR accelerated terminal differentiation of proerythroblasts (Figure 2D-F). These effects were prevented by adding TCDD to the culture, showing that SR1 directly targets AHR signaling to promote terminal differentiation of erythrocytes. We gained similar results from hESC-derived early erythroblasts and co-D14 CD34^+^ cell-derived proerythroblasts. Thus, our findings show that the role of AHR in erythroid differentiation is dependent upon the stage of erythroid differentiation. Initially, antagonizing AHR signaling promotes the generation of hPSC-derived HSPC, resulting in a parallel increased production of erythroid cells. While at early erythroid-megakaryocyte precursor stage, AHR signaling helps generate more erythroid cells before commitment to the erythroid lineage, but has no effect after that. Once committed to erythroid lineage, antagonism of AHR signaling enhanced the terminal differentiation of erythrocytes, irrespective of the origin of these erythrocytes, suggesting that suppression of AHR signaling conveys a regulatory mechanism on erythrocyte terminal differentiation and functional maturation that is common to all erythrocytes.

Through transcriptomic analysis by RNA-sequencing, we found that SR1 upregulated the expression of 162 overlapping genes in both UCB- and hESC-derived proerythroblasts. These genes were highly enriched in erythrocyte maturation processes, such as the erythrocyte differentiation and homeostasis, and heme metabolic processes. This finding demonstrated that antagonizing AHR signals comprehensively promotes the terminal differentiation and maturation of erythrocytes (Fig 3). Interestingly, RNA-seq results showed that SR1 downregulated the expression of genes highly associated with AHR (Figure S4A). Among these genes, the expression of *CYP1B1* (a gene that is downstream of AHR), but not *CYP1A1,* was downregulated. Since previous reports showed that *CYP1A1* is predominantly expressed in blood cells^50, 51^, our finding might highlight a novel role for *CYP1B1* in AHR-mediated oxidative stress during erythroid cell differentiation.

Further analysis revealed that SR1 treatment downregulated the expression of 440 genes common to both UCB- and hESC-derived proerythroblasts. In particular, actin cytoskeleton-related genes were highly enriched. Our findings suggest AHR antagonism promotes erythrocyte differentiation by modulating actin activity. Actin is an erythroid cytoskeleton protein that has an essential role in normal erythropoiesis^32^, and so our finding may be important in better understanding the mechanism by which AHR regulation promotes the terminal maturation of erythrocytes. During terminal erythroid differentiation, the expression of actin peaks in proerythroblasts and then decreases in subsequent differentiation stages^52^. In our experiments, we found that SR1-treated cells had lower F-actin expression than TCDD-treated cells, in which F-actin expression did not alter at D3 (Figure 4C). SR1 treatment accelerated F-actin remodeling and caused F-actin to be localized to one side of the cell near the constricted nucleus. This finding is in agreement with reports showing that during erythroid cell maturation, F-actin rearranges and forms a contractile ring between the extruding nucleus and the incipient reticulocyte; this process is essential to enucleation^32^. Moreover, the effect of AHR antagonists on terminal erythroid differentiation was reversed by actin-disrupting drugs (Figure 4D-G). Thus, our results demonstrate that antagonizing AHR signaling promotes actin rearrangement during terminal erythroid differentiation. Taken together, these data show that antagonizing the AHR pathway can positively modulate actin-mediated cytoskeleton rearrangements and hemoglobin synthesis.

We, and other groups, have reported that a large number of functionally mature erythrocytes could be generated from hESC *in vitro*^41, 42, 53^, but it is still difficult to undergo enucleation for these cells. In our experiments on mature hESC-derived erythrocytes, inhibition of AHR signaling also promoted terminal erythroid differentiation, with cells showing a more mature morphology and better deformability. Moreover, SR1 treatment resulted in a 2-fold increase in the number of enucleated erythrocytes (Figure 5). This positive effect of AHR antagonists on erythropoiesis, coupled with SR1-facilitated production of hESC-derived HSPC, points to a novel approach for the large-scale production of mature erythroid cells from hPSC.

In summary, our study shows that antagonizing AHR signaling accelerates erythroid terminal differentiation. Our exploration of the mechanism that underlies this effect suggests that inhibition of the AHR pathway induces an oxidative stress response in proerythroblasts and accelerates the actin-mediated rearrangement of the cytoskeleton. Future studies aimed at better understanding the regulation of AHR signaling during terminal erythrocyte differentiation might uncover further molecular mechanisms. Our study also provides a novel approach for the large-scale production of hPSC-derived mature erythrocytes.

## Materials and Methods

### Maintenance of hESC

H1 hESCs were provided by Professor Tao Cheng. H1 hESCs were maintained as described previously^40, 41^.

### Differentiation of hESC into HSPCs

hESCs were differentiated into HSPCs as described previously^40, 41^. SR1 (750 nM) and DMSO (used as vehicle control) were added at co-culture day 6 (co-D6).

### Differentiation of HSPCs into erythroid cells

To generate erythroid cells from co-cultured cells, co-D12 or –D14 CD34^+^ cells were isolated using a human CD34 positive selection kit (Stem Cell Technologies, Vancouver, Canada). CD34^+^ cells (1 × 10^5^/mL) were suspended in serum-free expansion medium II (Stem Cell Technologies) with 50 ng/mL stem cell factor (PeproTech, Rocky Hill, NJ, USA), thrombopoietin (PeproTech), Fms-related tyrosine kinase 3 ligand (PeproTech), and 0.5% penicillin/streptomycin (Invitrogen, Carlsbad, CA, USA). After 3 days in HSPC expansion culture, the cells were cultured using a three-phase liquid culture system (described in the **Supplemental Methods**). To generate umbilical cord blood (UCB)-derived erythroid cells, CD34^+^ cells were sorted using the same method described above, and then the cells were cultured in a three-phase liquid culture system.

### Hematopoietic colony assay

Cells were culture on colony assay medium which were described previously^40–43^. BFU-E, CFU-E, CFU-GEMM, and CFU-G/M/GM colonies were assessed after 12-14 days.

### Flow cytometry and cell sorting

Co-culture cells were harvested as described previously^40^. Flow cytometry was performed by using FACS Canto II system (BD Biosciences, San Jose, CA, USA) and cells were sorted by using a FACS Cytometer System (FACSJazz, BD, Franklin Lakes, NJ, USA). The antibodies used are presented in Table S1. Data were analyzed using FlowJo software (v.10.0.8)

### Morphological observation and immunofluorescence staining analysis

Harvested cells were centrifuged on glass slides by a cell cytospin machine (Cytospin 4, Thermo Fisher Scientific, Waltham, MA, USA). Morphology and membrane proteins were observed by MGG and IF staining respectively as described previously^40, 41^. The antibodies used are presented in Table S1.

### RNA-sequencing (RNA-seq) and data analysis

UCB-derived and hESC-derived proerythroblasts (CD71^+^GPA^low^) treated with SR1 or DMSO were collected at D3. RNA-seq was performed on these cells by Novogene Co., Ltd. (Beijing, China). Gene ontology enrichment (EGO) analysis, Gene set enrichment analysis (GSEA) and heatmap analysis were performed on raw data as described previously^43^. The sequences data reported in this study were archived in the Sequence Read Archive with the accession number PRJNA724809.

### F-actin staining assay

Erythroid cells were harvested and washed three times with PBS. Cells were fixed with 4% paraformaldehyde for 10 min at room temperature, followed by permeabilization in 0.1% Triton X-100 for 2 min at room temperature. After washing, cells were blocked with 1% BSA for 20 min at room temperature, and then washed three times with PBS. Subsequently, fluorescent FITC-conjugated phalloidin (MedChem Express, Monmouth Junction, NJ, USA) diluted in 1% BSA (5 μg/mL) was added and incubated for 30 min at 37°C. After nuclear staining with Hoechst 33342 (Invitrogen), cells were centrifuged on glass slides. Images were acquired using a ZEISS LSM900 Airyscan microscope.

### Deformability measurements

Erythroid cells (1 × 10^6^) were re-suspended in 10μl PBS and added to 1 mL polyvinylpyrrolidone solution. The total cell suspension was transferred to erythrocyte deformability tester (LBY-BY, Beijing, China) to test the deformation of erythroid cells under shear-rate conditions of 1000 s^-1^.

### Statistical analysis

All data are shown as the mean ± SD from three independent experiments. Statistical significance was evaluated using the Student’s t-test (when two groups were analyzed) or one-way ANOVA with Tukey’s multiple testing correction as appropriate (for three or more groups). A p-value < 0.05 was considered significant.

## Acknowledgments

We thank Professor Tao Cheng at State Key Laboratory of Experimental Hematology, Institute of Hematology and Blood Diseases Hospital, CAMS & PIMC for generously providing the H1 line. We also thank Professor Duonan Yu for precious suggestions on this work. This work was supported by the National Basic Research Program (973 Program; 2015CB964902), the National Natural Science Foundation of China (H81170466 and H81370597), the CAMS Initiatives for Innovative Medicine (2016-I2M-1-018, 2019-I2M-1-006, 2017-I2M-2005) and the National Nature Science Foundation of China Youth Fund (81700107) to B.M..

## Author contributions

Y.J.C. performed research, analyzed data; Y.D. performed RNA-seq and proteomics data analysis. X.L.L., W.J.L., Y.M.Z., B.M., X.P., X.H.L., Y.Z., Q.M.A., F.X.X., Y.X., X.P.C, M.W.L., Q.X.Z., and Y.G.Z. performed some of the experiments. Y.J.C. and F.M. designed the project, discussed the data, and wrote the manuscript. F.M. approved the final manuscript.

## Supplemental methods

### Erythroid differentiation in three-phase liquid culture system

In the first phase (day0-7), cells were culture in SFEM II supplemented with 10^-6^ M dexamethasone (Sigma-Aldrich, St. Louis, MO, USA), 0.5% penicillin/streptomycin, 100 ng/ml SCF, 5 ng/ml interleukin-3 (PeproTech, Rocky Hill, NJ, USA), 4 U/ml erythropoietin (PeproTech). In phase 2 (day 7-12), both Dex and IL-3 were omitted from the medium. In phase 3 (day12-17), cells were cultured at a density of 10^6^ cells/ml, with DEX, SCF and IL-3 omitted from the medium and supplementing 1 mg/ml holo-transferrin (Sigma-Aldrich), 20% fetal bovine serum (Biological Industries, Kibbutz Beit Haemek, Israel), and 10% human AB serum (Gemini, Woodland, CA, USA).

### Isolation and culture of hematopoietic endothelial cells and hematopoietic progenitor cells

Co-D6 CD34^+^KDR^+^GPA^-^ and co-D10 CD34^+^CD43^+^GPA^-^ cells were sorted and re-plated on 10^4^ AGM-S3 cells in hematopoietic differentiation medium which was described previously.

### Apoptosis analysis

Cells were stained with annexin V-APC and 7-AAD according to the manufacture’s instructions (Biolegend, San Diego, CA, USA) and examined by flow cytometry.

### Cell-cycle analysis

Cells were treated with BrdU for 8 hours, and processed using an APC-BrdU Flow Kit (BD Biosciences, San Jose, CA, USA) according to the manufacturer’s instructions.

### Proteomics and data analysis

UCB-derived proerythroblasts with SR1 or DMSO dealing were collected at D3. Proteomics using the MaxLFQ label-free quantification was performed by Shanghai Apploed Protein Technology Co., Ltd. (Shanghai, China). Normalized data were gained and heatmap was plotted as described previously. Data are available via ProteomeXchange with identifier PXD025624.

### Western blot

Proteins were separated by SDS-PAGE and transferred to PVDF (Merck Millipore, Billerica, MA, USA). After 5% milk blocking for 1 hour, membranes were incubated with primary antibody (mouse anti-human β-actin; mouse anti-human GAPDH; all Beyotime, Shanghai, China) at 1:5000 dilution for overnight at 4 ℃. Membranes were incubated with Second antibody (goat anti-mouse immunoglobulin-HRP, Beyotime) at 1:2000 dilution for 1 hour at RT. Bands were visualized by enhanced chemilunescence (GE Healthcare, Little Chalfont, Buckinghamshire, UK) and exposed to Imagequant Las 4000 (USA).

---

**Supplemental table 1.** Antibodies used in this study

| Antibodies | Company |
| --- | --- |
| PE-conjugated hemoglobin $\beta$ | Santa Cruz Biotechnology (Santa Cruz ,CA, USA) |
| PE-conjugated CD47 | BD Pharmingen (San Diego, CA, USA) |
| PE-conjugated CD44 | BD Pharmingen |
| PE-conjugated CD49d | BD Pharmingen |
| PE-conjugated AHR | eBioscience (San Diego, CA, USA) |
| PE-conjugated CD29 | BD Pharmingen |
| Hemoglobin | Bethyl Laboratories (Montgomery, TX, USA) |
| hemoglobin $\beta$ | Santa Cruz Biotechnology |
| Donkey anti-goat IgG (H+L) Cy3 | Jackson Immuno Research<br>(West Grove, PA, USA) |
| Donkey anti-goat IgG (H+L) FITC | Jackson Immuno Research |
| Hoechst 33342 | Invitrogen (Carlsbad, CA, USA) |
| SYTO16 | Invitrogen |

## Supplemental Figures

**Supplemental Figure 1.**
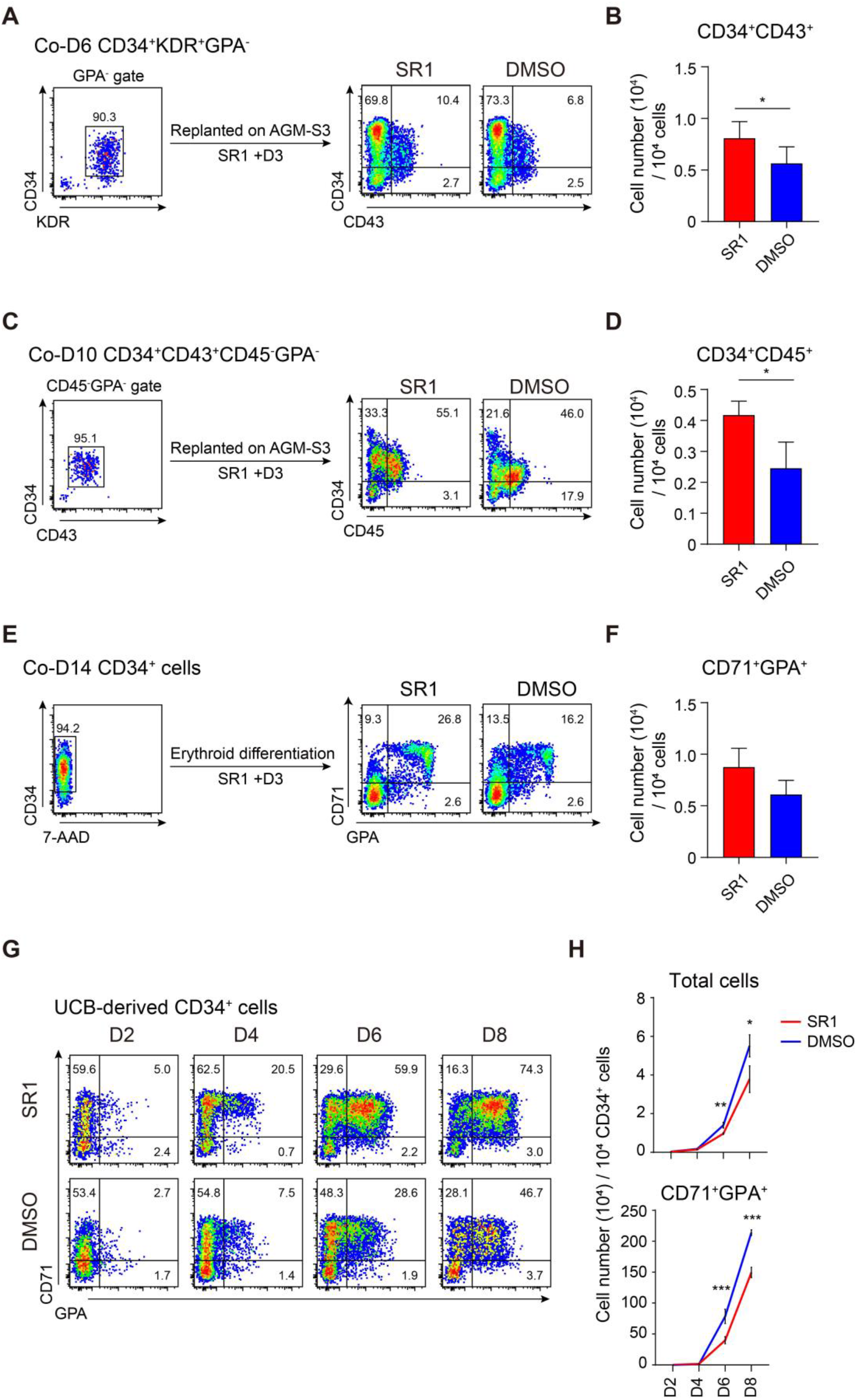
SR1 do not promote erythroid cell generated from hematopoietic progenitor. Isolated co-D6 hematopoietic endothelial cells (HEC) and Co-D10 hematopoietic progenitor cells (HPC) were replanted on AGM-S3 containing hematopoietic differentiation medium with or without SR1. The purity of HEC and HPC is shown; the proportion of HEC-derived CD34^+^CD43^+^ (A) and HPC-derived CD34^+^CD45^+^ (C) cells was analyzed by flow cytometry at D3. The absolute number of HEC-derived CD34^+^CD43^+^ (B) and HPC-derived CD34^+^CD45^+^ (D) cells was calculated at D3. Isolated co-D14 CD34^+^ cells were re-cultured in erythroid differentiation medium. The purity of CD34^+^ cells is shown, and the proportion of CD71^+^GPA^+^ cells (E) was analyzed by flow cytometry at D3. The absolute number of CD71^+^GPA^+^ cells (F) was calculated at day 3. Isolated UCB-derived CD34+ cells were cultured in erythroid differentiation medium. The proportion of CD71^+^GPA^+^ cells (G) was analyzed by flow cytometry and the total number of cells and CD71^+^GPA^+^ cells (H) was calculated at days 2, 4, 6, and 8. All values are mean ± SD, N = 3; *p < 0.05, ** p < 0.01, *** p < 0.001.

**Supplemental Figure 2.**
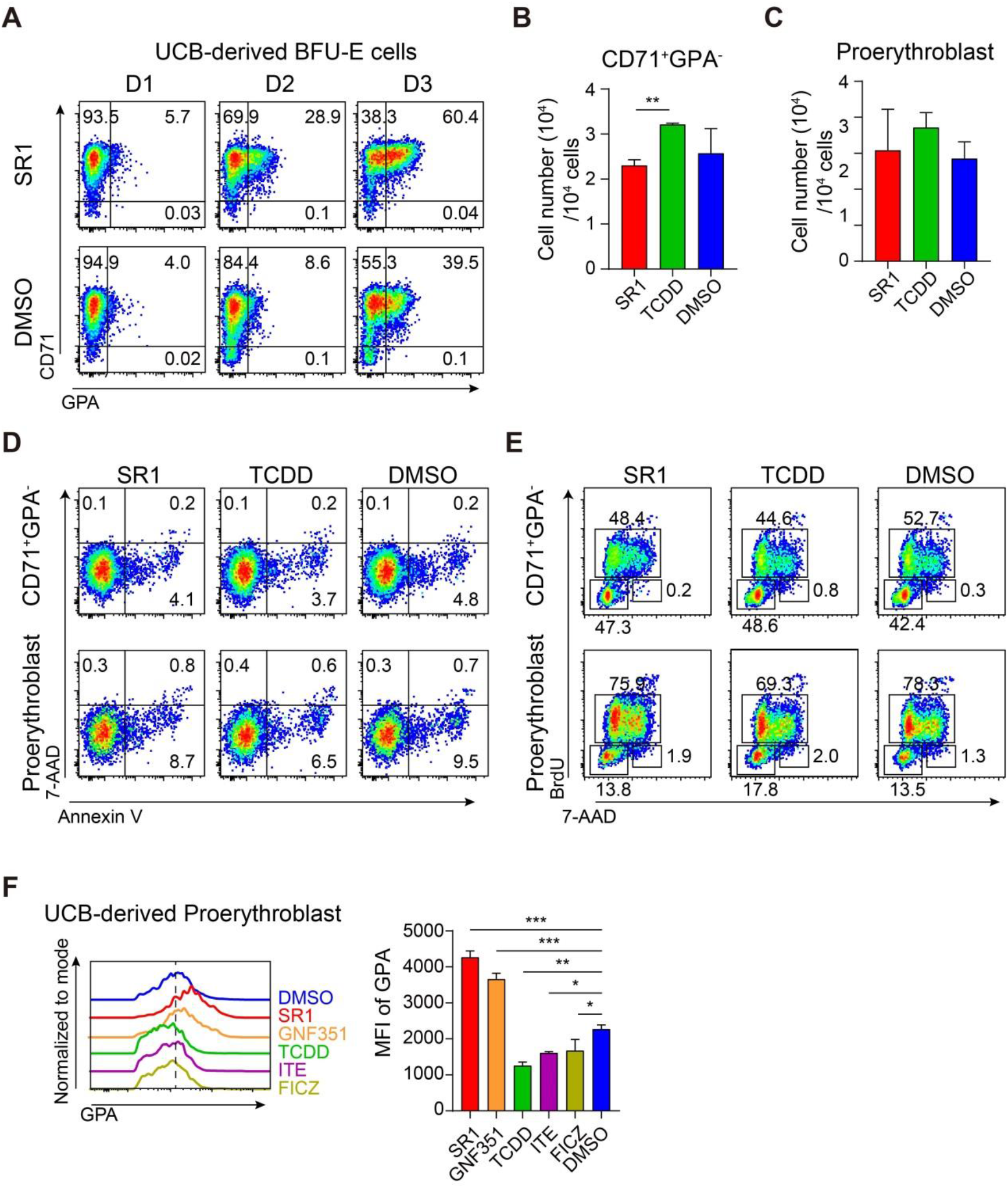
AHR Inhibition improves erythroid terminal differentiation without influencing cell cycle and apoptosis. Sorted UCB-derived BFU-E cells were continuously treated with or without SR1. (A) CD71 and GPA expression was analyzed by flow cytometry at days 1, 2, and 3. Isolated UCB-derived proerythroblasts were treated with or without SR1, the total number of cells derived from CD71^+^GPA^-^ cells (B) and proerythroblasts (C) was calculated at D3. Apoptosis (D) and the cell cycle (E) of CD71^+^GPA^-^ cells and proerythroblasts were analyzed by flow cytometry at D3. Sorted proerythroblasts cells were exposed to SR1, GNF351, TCDD, ITE, FICZ, or DMSO; (F) the expression of GPA was analyzed by flow cytometry, and the mean fluorescence intensity (MFI) of GPA was calculated at D3.

**Supplemental Figure 3.**
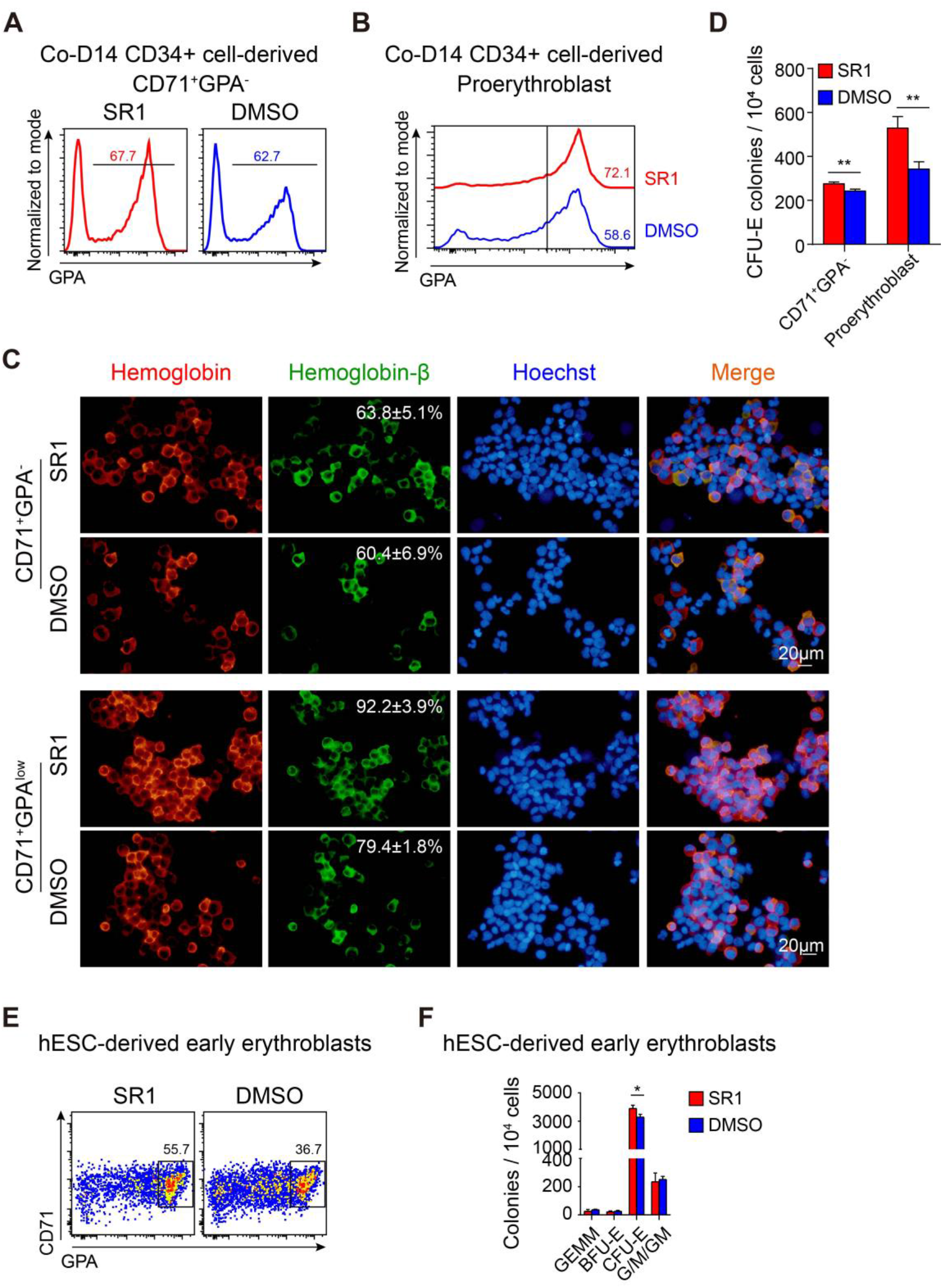
SR1 promotes H1-derived erythroid terminal differentiation. H1-derived CD71^+^GPA^-^ cells and proerythroblasts were treated with or without SR1. The proportion of GPA^+^ cells generated from CD71^+^GPA^-^ cells (A) and GPA expression in proerythroblasts (B) was analyzed by flow cytometry at D3. Hemoglobin-β expression was detected by IF staining, and the percentage of β-positive cells was calculated at day 3. (C) Representative images of stained cells are shown; hemoglobin, red; hemoglobinβ, green; Hoechst 33342 (nuclei), blue; scale bar, 20μm. (D) The clonogenic capacity of H1-derived CD71^+^GPA^-^ cells and proerythroblasts treated with or without SR1 at D3. The number of CFU-E colonies is shown. H1-derived early erythroblasts were sorted and exposed to SR1 or DMSO for 3 days. (E) The proportion of CD71^+^GPA^high^ cells was analyzed by flow cytometry at D3. (F) The clonogenic capacity of H1-derived early erythroblasts. All values are mean ± SD, N = 3; * p < 0.05, ** p < 0.01, *** p < 0.001.

**Supplemental Figure 4.**
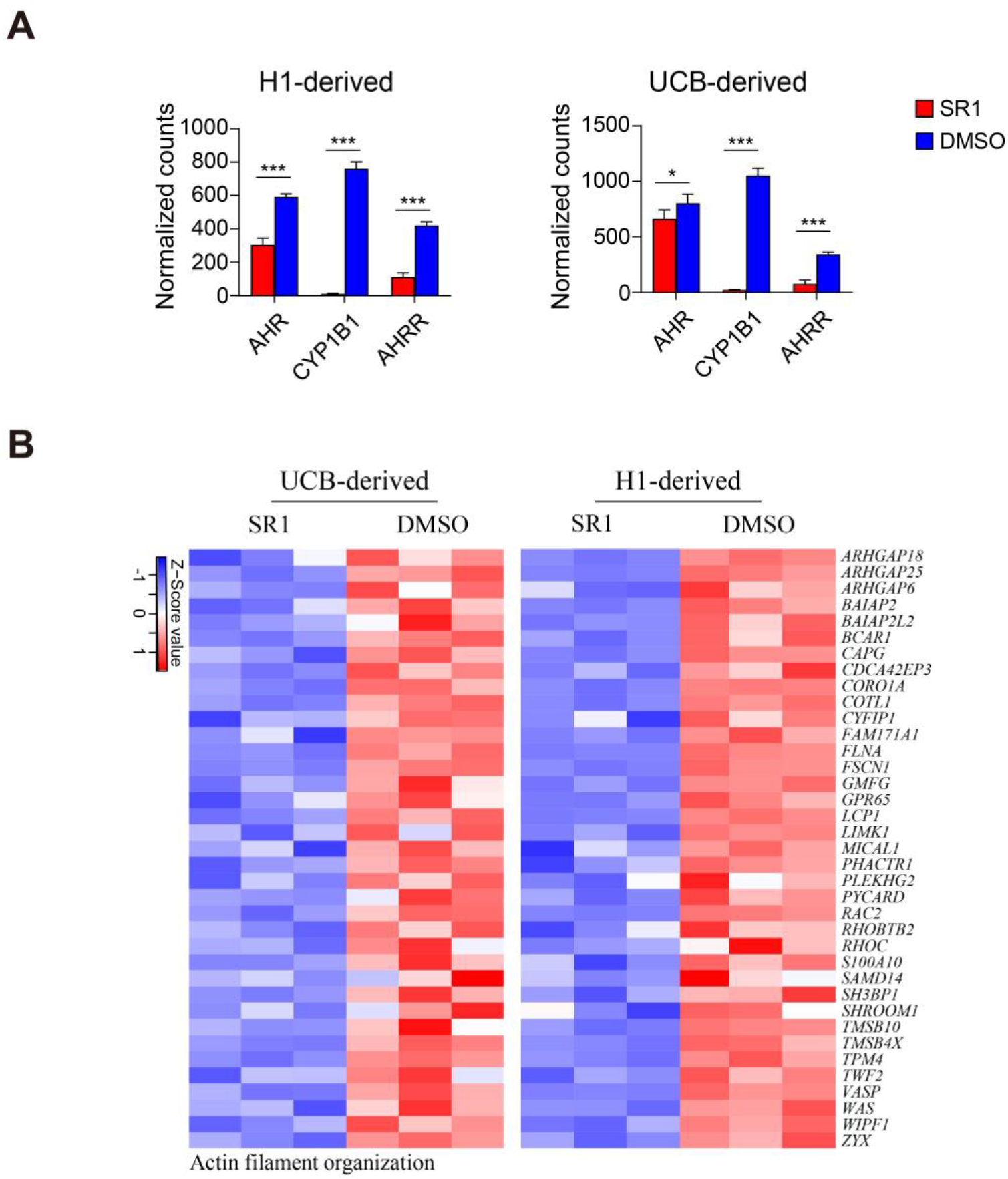
SR1 regulated AHR- and F-actin-related gene. UCB- and H1-derived proerythroblasts was performed RNA-seq following treatment with or without SR1 at days 3. (A) The expression of *AHR*, *CYP1B1* and *AHRR* in H1- and UCB-derived proerythroblasts from RNA-seq data. (B) Heatmaps representing actin filament organization-related genes identified from the results of ontology enrichment analysis in UCB-derived and H1-derived proerythroblasts; columns represent the indicated replicates of each population; the color bar shows row-standardized Z-scores.

**Supplemental Figure 5.**
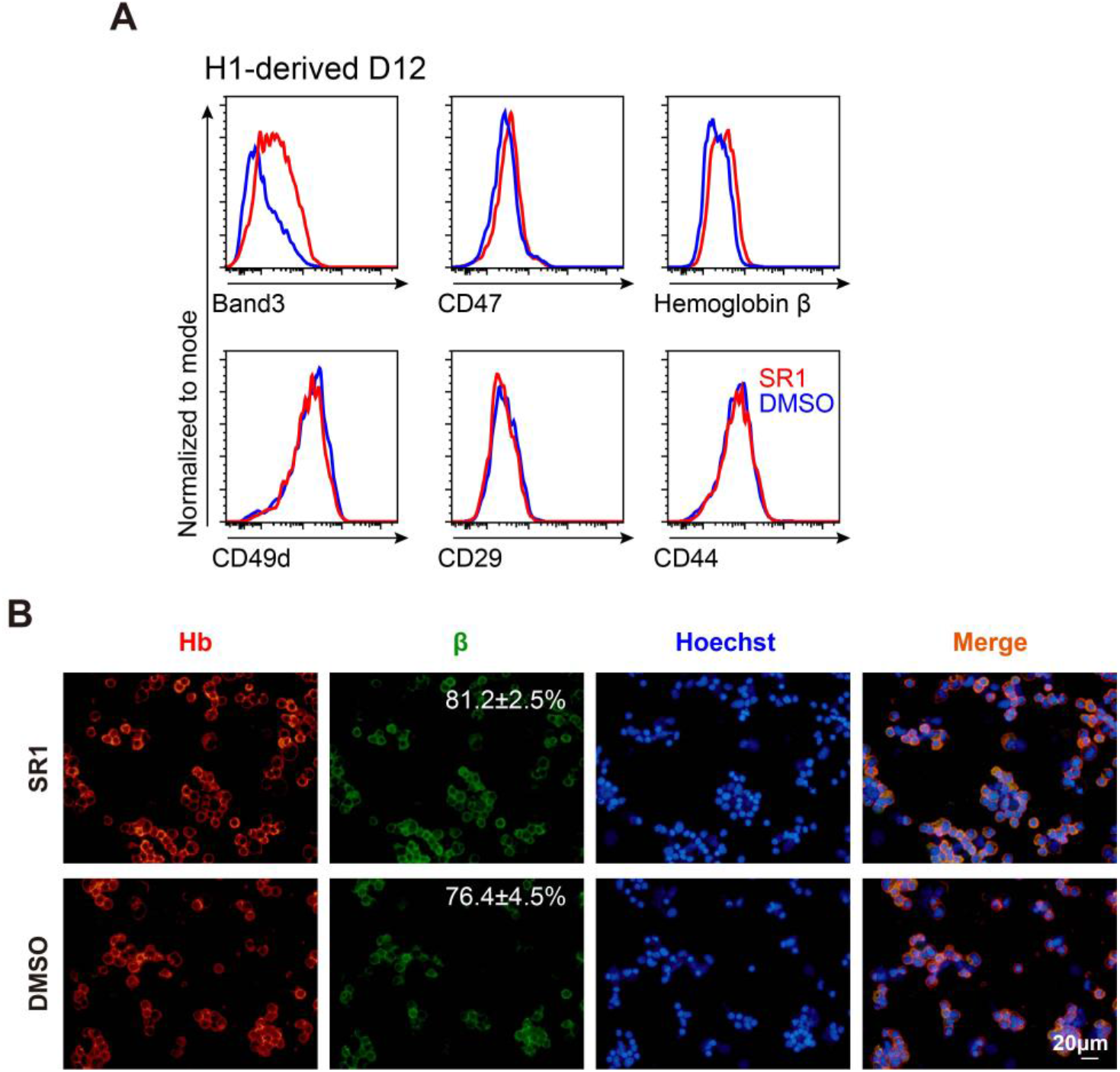
SR1 enhances H1-derived erythroid cells maturation. SR1 was added on D7 during H1-derived erythroid differentiation, and erythroid cells were collected at D12. (A) The expression of erythroid-related proteins was analyzed by flow cytometry at day 12. (B) Representative images of stained cells are shown; hemoglobin, red; hemoglobinβ, green; Hoechst 33342 (nuclei), blue; scale bar, 20μm.

## Notes

### Competing Interest Statement

The authors have declared no competing interest.

